# Signal bias at glucagon family receptors: rationale and downstream impacts

**DOI:** 10.1101/2020.04.26.062372

**Authors:** Ben Jones, Emma Rose McGlone, Zijian Fang, Phil Pickford, Ivan R Corrêa, Atsuro Oishi, Ralf Jockers, Asuka Inoue, Sunil Kumar, Frederik Görlitz, Chris Dunsby, Paul MW French, Guy A Rutter, Tricia Tan, Alejandra Tomas, Stephen R Bloom

## Abstract

Receptors for the peptide hormones glucagon-like peptide-1 (GLP-1), glucose-dependent insulinotropic polypeptide (GIP) and glucagon (GCG) are important regulators of insulin secretion and energy metabolism. Recently described GLP-1 receptor agonists showing signal bias in favour of cyclic AMP over β-arrestin-2 recruitment have delivered promising results in preclinical studies. Here we first sought to establish the role of β-arrestins in the control of intracellular signalling and trafficking responses at the closely related GLP-1, GIP and GCG receptors, through studies performed in cells depleted of both β-arrestin isoforms. We also generated analogues of GLP-1, GCG and GIP which in some cases showed selective reduction in β-arrestin-2 recruitment *versus* cAMP signalling compared to the parent peptide. Despite reduced acute signalling potency and/or efficacy, some biased GLP-1 and GIP analogues increased maximal sustained insulin secretion from INS-1 832/3 clonal beta cells, although only at high agonist concentrations. Biased GCG analogues did not affect maximal insulin release, or glucose output in hepatocytes.

## Introduction

The receptors for the glucagon-like peptide-1 (GLP-1R), glucose-dependent insulinotropic polypeptide (GIPR) and glucagon (GCGR) are major pharmacological targets in metabolic diseases such as type 2 diabetes (T2D) and obesity (1). Each of these receptors is present on pancreatic beta cells, and an important component of their overall metabolic actions when physiologically or pharmacologically activated is augmentation of glucose-stimulated insulin release (2, 3). In hepatocytes, GCGR facilitates glucose output, which may be undesirable in T2D; however, its “energy wasting” effect in peripheral tissues (2) could mitigate hyperglycaemia by weight loss and associated improvements in insulin sensitivity.

GLP-1R, GIPR and GCGR are closely related G protein-coupled receptors (GPCRs) of the class B (secretin) family. When activated, they engage the G protein Gα_s_, which is coupled to insulin secretion via generation of cyclic adenosine monophosphate (cAMP) (4), and β-arrestins, which are multifunctional scaffold proteins widely reported to initiate non-G protein signalling cascades such as phosphorylation of mitogen-activated protein kinases (MAPKs) (5), and concurrently diminish G protein signalling by steric hindrance (6). To varying degrees, each of these receptors undergoes agonist-mediated endocytosis (7), which fine tunes the spatial origin and duration of their intracellular signalling responses (8, 9).

The balance between recruitment and activation of intracellular signalling effectors and subsequent receptor trafficking can be ligand-specific – a pharmacological concept known as “biased signalling” (10). A number of examples of signal bias at the GLP-1R have been described, including both naturally occurring (11, 12) and pharmacological (13, 14) orthosteric agonists. Importantly, G protein-biased GLP-1R agonists derived from exendin-4 lead to increases in sustained insulin secretion through avoidance of GLP-1R desensitisation, reduction of GLP-1R endocytosis and resultant attenuation of GLP-1R downregulation over pharmacologically relevant time periods (15, 16). Moreover, a GLP-1R/GIPR dual agonist (Tirzepatide) with promising results for the treatment of T2D in clinical trials (17) has recently been reported to show pronounced G protein bias at the GLP-1R, although not at the GIPR (18). In view of the current drive to develop incretin analogues jointly targeting GLP-1R, GCGR and GIPR (1), we sought to establish whether signal bias could similarly be achieved at the latter two receptors, and to determine if this is associated with prolonged signalling responses, as seen with the GLP-1R.

In this work, we first compare β-arrestin recruitment and activation profiles of GLP-1R, GIPR and GCGR activated by their cognate ligands, and subsequently demonstrate how the absence of β-arrestins affects patterns of intracellular signalling and trafficking. We also find that a number of substitutions close to the N-terminus of the cognate ligand for each receptor result in reductions in both cAMP signalling and β-arrestin-2 recruitment, with bias in favour of cAMP in some cases. However, compared to our previous study with biased exendin-4 analogues at the GLP-1R (15), the degree of bias achieved here was more modest. Moreover, whilst bias-related differences were apparent in downstream responses such as insulin secretion, these only occurred at high agonist concentrations.

## Results

### Coupling of GLP-1R, GIPR and GCGR to intracellular effectors and endocytosis

We first performed studies to compare responses to the cognate agonist for each receptor in HEK293T cells. Specifically, for GLP-1R we used GLP-1(7–36)NH_2_, for GIPR we used GIP(1–42), and for GCGR we used full length GCG(1–29). These ligands are referred to henceforth as GLP-1, GIP and GCG. Using NanoBiT complementation (19) to detect ligand-induced recruitment of LgBiT-tagged mini-G proteins (20) to each of the receptors tagged at the C-terminus with the complementary SmBiT sequence, we confirmed a robust ligand-induced mini-G_s_ response, but more minor increases with mini-G_q_ and mini-G_i_ (Figure 1A, Supplementary Figure 1A). This is in keeping with the consensus that glucagon family receptors are primarily coupled to cAMP signalling via Gα_s_, with system-dependent engagement with other Gα sub-types under some circumstances (21, 22). Moreover, LgBiT-β-arrestin-2 recruitment responses could be detected in all cases but were more transient than for mini-G proteins, matching the pattern seen with pharmacological GLP-1R agonists (23). Notable differences between receptor types included 1) substantially greater amplitude for mini-G_s_ recruitment for GLP-1R than for GCGR and GIPR, with the latter also showing slower kinetics (t_1/2_ = 7.1 ± 0.4 min versus 1.5 ± 0.2 min for GLP-1R, p<0.05 by unpaired t-test); 2) mini-G_i_ and mini-G_q_ responses were virtually undetectable for GIPR; and 3) markedly reduced recruitment of β-arrestin-2 to GIPR compared to GLP-1R and GCGR, in keeping with another report (24). These and other responses are also quantified from their AUC in Figure 1G as well as Supplementary Figure 1A.

**Figure 1.**
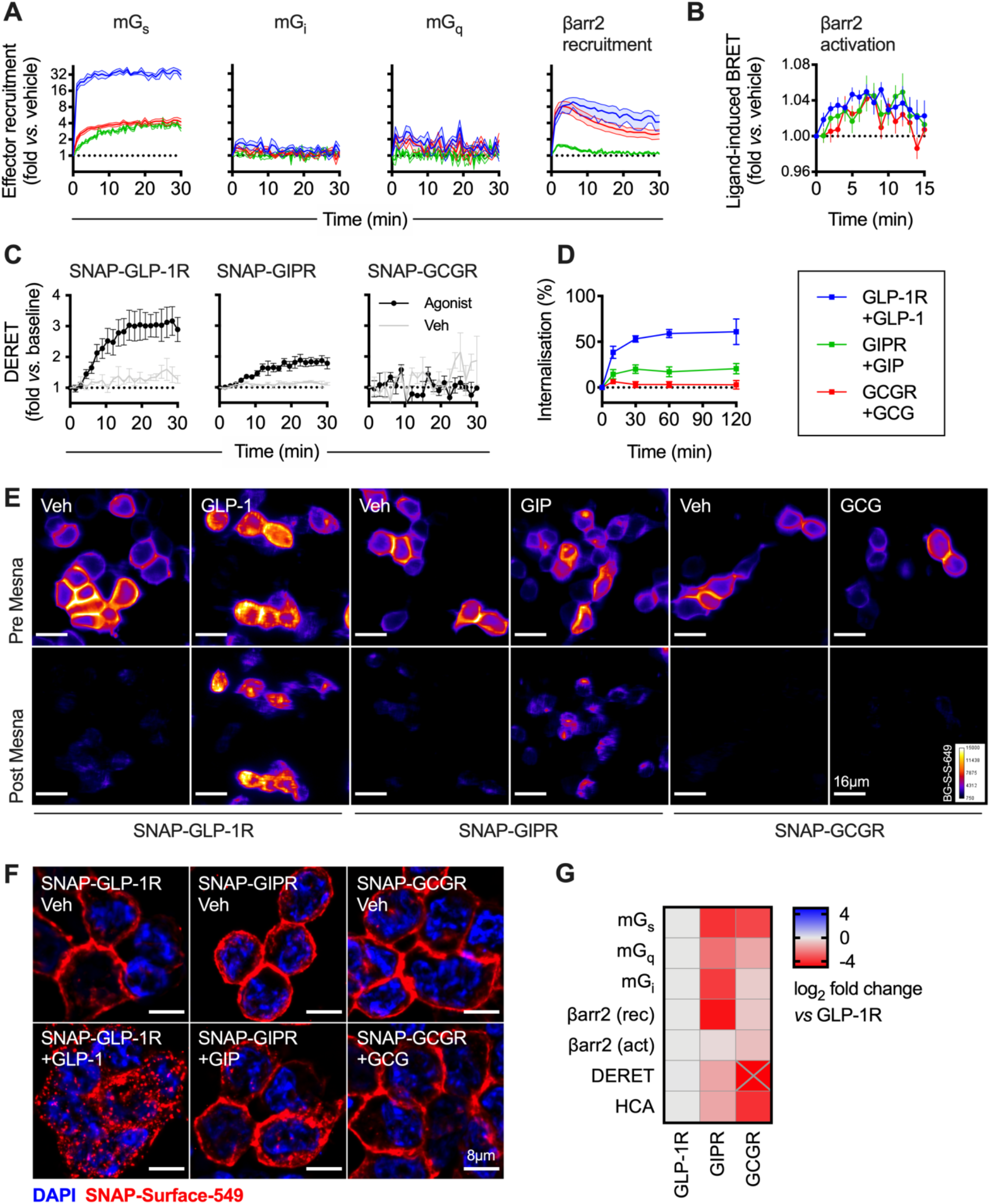
Signalling and internalisation properties of GLP-1R, GIPR, GCGR. (**A**) Mini-G_s_, -G_i_, -G_q_, and β-arrestin-2 recruitment responses in HEK293T cells transiently transfected with each SmBiT-tagged receptor and LgBiT-tagged effector and stimulated with 100 nM GLP-1, GIP, GCG or vehicle; *n*=4 for GLP-1R and GCGR and *n*=5 for GIPR. (**B**) β-arrestin-2 activation response in HEK293T cells transiently transfected with each receptor and NLuc-4myc-βarr2-CyOFP1 and stimulated with 100 nM GLP-1, GIP, GCG or vehicle, *n*=6. (**C**) Internalisation of each SNAP-tagged receptor in HEK293T cells stimulated with 100 nM GLP-1, GIP, GCG or vehicle, measured by DERET, *n*=4. (**D**) Time course showing internalisation of each SNAP-tagged receptor in response to 100 nM GLP-1, GIP or GCG, measured by reversible SNAP-tag labelling in transiently transfected HEK293T cells, *n*=5. (**E**) Representative images of transiently transfected HEK293T cells expressing each SNAP-tagged receptor and labelled with BG-S-S-649 prior to treatment ± 100 nM GLP-1, GIP or GCG. Images were acquired before and after removal of residual surface BG-S-S-649 using Mesna; scale bar = 16 μm. (**F**) Representative high-resolution images from *n*=2 experiments of HEK293T cells transiently expressing each SNAP-tagged receptor and labelled with SNAP-Surface-549 before treatment ± 100 nM GLP-1, GIP or GCG; scale bar = 8 μm. (**G**) Heatmap summary of signalling and internalisation responses shown in this figure, normalised to the GLP-1R response and expressed as a log_2_ fold change. β-arrestin-2 recruitment (“rec” – Figure 1A) and activation (“act” – Figure 1B) responses are both shown. GCGR DERET response falls below the displayed range and is marked with “X”. See also Supplementary Figure 1. Data are represented as mean ± SEM.

After recruitment to activated GPCRs, β-arrestins undergo conformational rearrangements which are important for their functions (25, 26). Using a recently developed intramolecular BRET-based biosensor (27) we compared the ability of GLP-1R, GIPR and GCGR to activate β-arrestin-2 when stimulated by their cognate agonists. Here, comparable ligand-induced BRET signals were detected with each receptor (Figure 1B, Supplementary Figure 1B), highlighting how measuring recruitment of intracellular effectors *per se* may not provide the full picture for how a receptor or ligand can engage different intracellular pathways. Note that the BRET ratio obtained in the presence of each transfected receptor prior to stimulation was identical, arguing against receptor-specific differences in constitutive β-arrestin-2 activation (Supplementary Figure 1B).

β-arrestin recruitment is classically linked to GPCR endocytosis (28), although conflicting evidence exists for its role in controlling trafficking of incretin receptors (29–33). We first used diffusion-enhanced resonance energy transfer (DERET) (34) to monitor agonist-induced loss of surface labelled SNAP-tagged receptors transiently expressed in HEK293T cells. Robust internalisation was noted for GLP-1R, whereas GIPR internalisation was less extensive, and no ligand-induced change in DERET signal could be detected for GCGR (Figure 1C, Supplementary Figure 1C). Endocytic profiles were confirmed using an alternative approach based on reversible SNAP-tag labelling, in which the fluorescent probe BG-S-S-649 is cleaved from residual surface receptors after agonist-induced internalisation using the cell-impermeant reducing agent Mesna (16, 35). A time-course study showed rapid and extensive loss of surface SNAP-GLP-1R after GLP-1 treatment, whereas internalisation of the other two class B GPCRs was more limited (GIPR) or virtually absent (GCGR) (Figure 1D, Supplementary Figure 1D). Examples of the effect of Mesna cleavage are shown in Figure 1E, and higher resolution images showing ligand-induced distribution changes of surface-labelled SNAP-GLP-1R, -GIPR and -GCGR are shown in Figure 1F.

Comparing the measured responses for each receptor with GLP-1R as the reference, two notable observations were that 1) the largest amplitude responses were seen with GLP-1R for all readouts, and 2) GCGR showed greater ligand-induced recruitment of β-arrestin-2 (and other effectors) than GIPR, yet this did not translate to a corresponding increase in ligand-induced endocytosis (Figure 1G).

### Effect of β-arrestin depletion on GLP-1R, GIPR and GCGR signalling and trafficking

To further investigate the role of β-arrestins in incretin receptor signalling we used HEK293 cells in which both β-arrestins were deleted by CRISPR/Cas9 (36). Firstly, cAMP signalling responses to each cognate ligand were compared in wild-type or β-arrestin-knockout cells transiently transfected with the relevant SNAP-tagged receptor. In wild-type cells, a robust cAMP response was observed after 10 minutes stimulation, but after 60 minutes, efficacy was reduced (Figure 2A, Table 1). In contrast, the reduction in efficacy over time was much less marked in β-arrestin-knockout cells, suggesting that β-arrestins do, as expected, contribute to attenuation of Gα_s_ responses for GLP-1R, GIPR and GCGR. We also used the FRET biosensor AKAR4-NES (37) to detect cytoplasmic protein kinase A (PKA) activation in each cell type (Supplementary Figure 2A). Plate-reader measurements at multiple agonist concentrations allowed construction of dose-response curves from the overall AUC (Figure 2B, Table 2), demonstrating that potency for PKA activation was increased in β-arrestin-knockout cells. This suggests that increased cAMP signalling in the absence of β-arrestins is also propagated to downstream targets.

**Figure 2.**
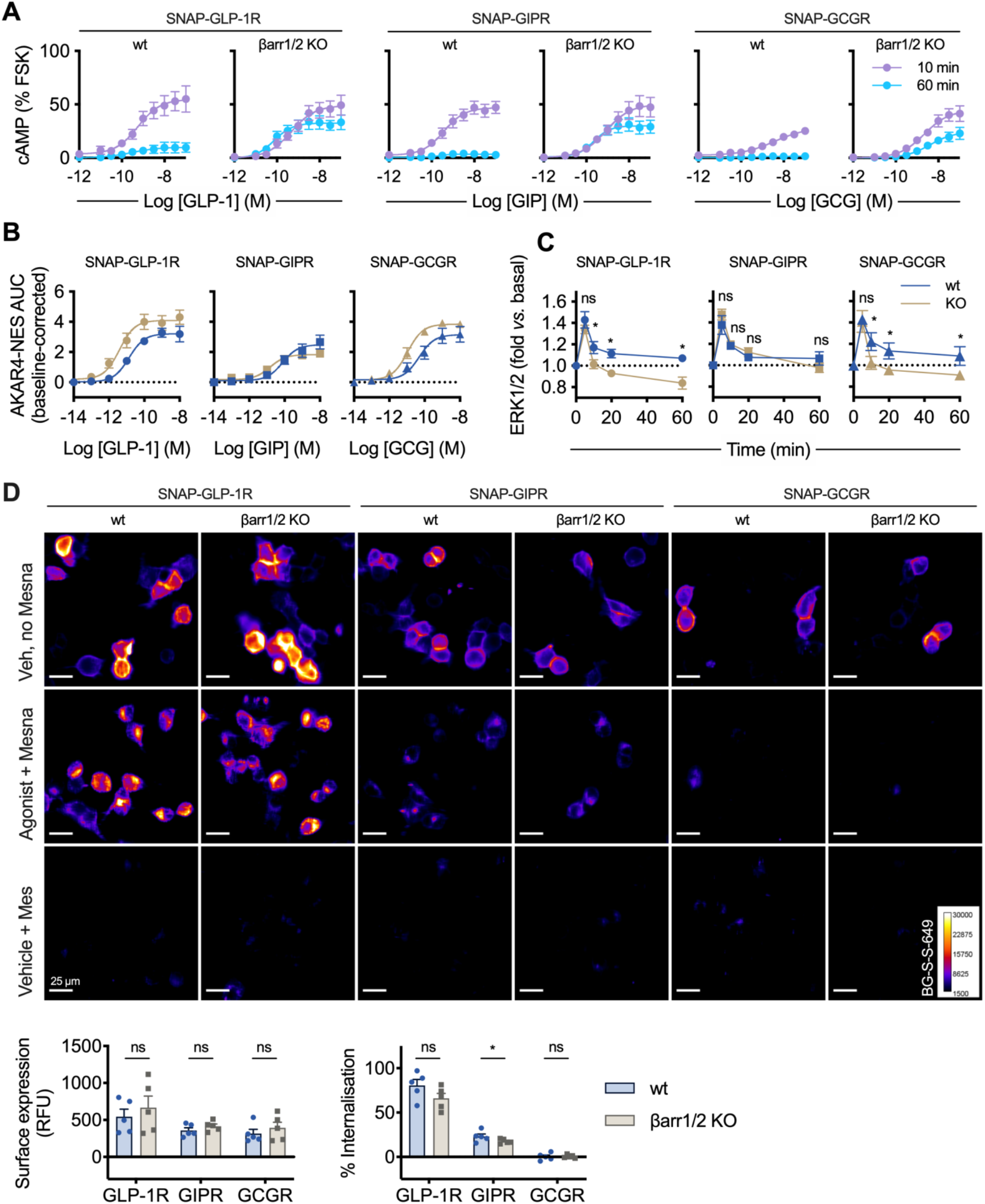
GLP-1R, GIPR and GCGR responses in the absence of β-arrestins. (**A**) cAMP responses at 10 or 60 min to GLP-1, GIP or GCG, in wild-type (wt) or dual β-arrestin knockout (βarr1/2 KO) HEK293 cells transiently transfected with the indicated SNAP-tagged receptor, normalised to forskolin (FSK, 10 μM) response, respectively, *n*=5, 4-parameter fits shown. (**B**) Dose responses for PKA activation in wild-type or dual β-arrestin knockout HEK293 cells transiently co-transfected with each SNAP-tagged receptor and AKAR4-NES, calculated from FRET signal AUC and normalised to vehicle responses, 3-parameter fits shown, *n*=5. See also Supplementary Figure 2. (**C**) Time-course for ERK1/2 phosphorylation in wild-type or dual β-arrestin knockout HEK293 cells transiently transfected with each SNAP-tagged receptor and stimulated with 100 nM GLP-1, GIP or GCG, normalised to basal response. Time-points compared by randomised block two-way ANOVA with Sidak’s test, *n*=6. (**D**) Representative images showing internalisation of each transiently transfected SNAP-tagged receptors in wild-type or dual β-arrestin knockout HEK293 cells, labelled with BG-S-S-649 prior to 100 nM agonist (or vehicle) treatment, with removal of residual surface receptor using Mesna where indicated; scale bar = 25 μm. Surface expression and ligand-induced internalisation are quantified from *n*=5 experiments, with comparisons by paired t-tests. * p<0.05 by statistical test indicated in the text. Data are represented as mean ± SEM with individual replicates in some places.

**Table 1.**
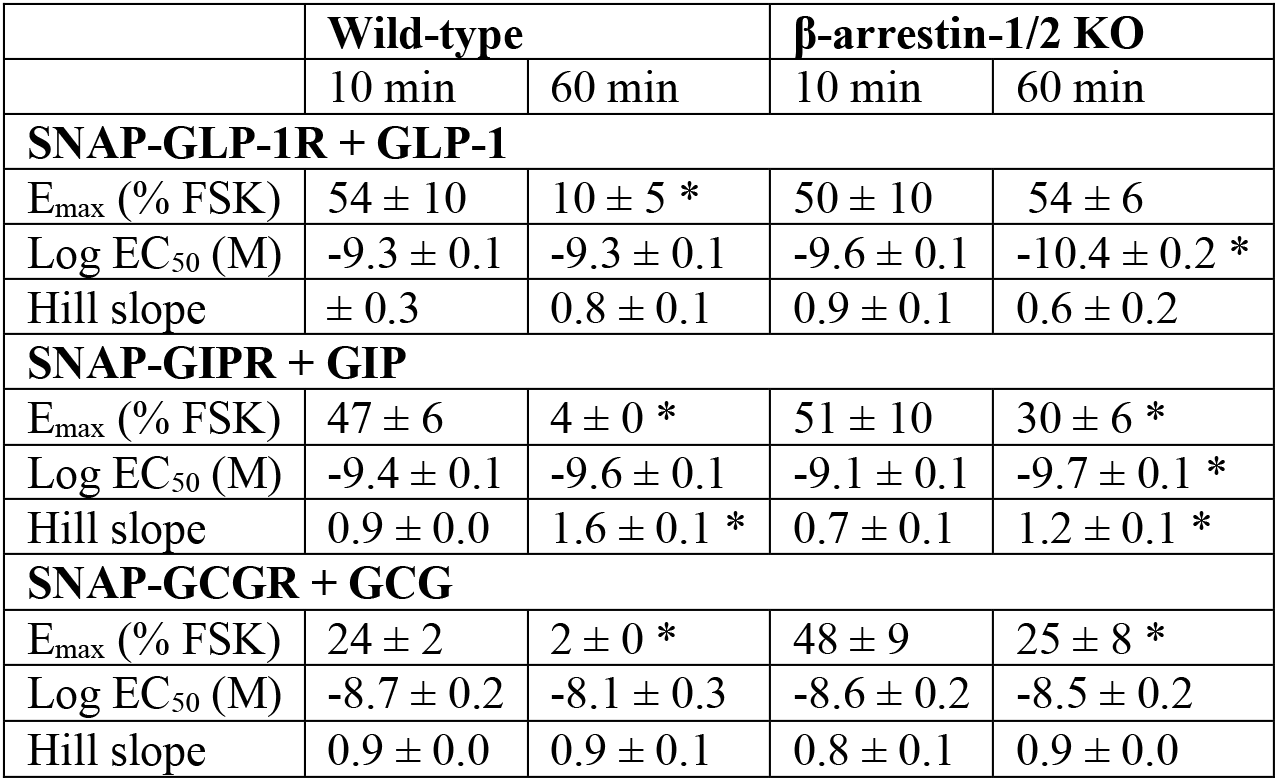
Effect of β-arrestin deletion on agonist-induced cAMP responses in HEK293 cells. Mean parameter estimates ± SEM from responses depicted in Figure 2A, *n*=5. * p<0.05 by two-way randomised block ANOVA with Sidak’s test comparing 10 min *versus* 60 min.

| | Wild-type | | β-arrestin-1/2 KO | |
| --- | --- | --- | --- | --- |
|  | 10 min | 60 min | 10 min | 60 min |
| **SNAP-GLP-1R + GLP-1** |  |  |  |  |
| $E_{\max}$ (% FSK) | $54 \pm 10$ | $10 \pm 5$ * | $50 \pm 10$ | $54 \pm 6$ |
| Log $EC_{50}$ (M) | $-9.3 \pm 0.1$ | $-9.3 \pm 0.1$ | $-9.6 \pm 0.1$ | $-10.4 \pm 0.2$ * |
| Hill slope | $\pm 0.3$ | $0.8 \pm 0.1$ | $0.9 \pm 0.1$ | $0.6 \pm 0.2$ |
| **SNAP-GIPR + GIP** |  |  |  |  |
| $E_{\max}$ (% FSK) | $47 \pm 6$ | $4 \pm 0$ * | $51 \pm 10$ | $30 \pm 6$ * |
| Log $EC_{50}$ (M) | $-9.4 \pm 0.1$ | $-9.6 \pm 0.1$ | $-9.1 \pm 0.1$ | $-9.7 \pm 0.1$ * |
| Hill slope | $0.9 \pm 0.0$ | $1.6 \pm 0.1$ * | $0.7 \pm 0.1$ | $1.2 \pm 0.1$ * |
| **SNAP-GCGR + GCG** |  |  |  |  |
| $E_{\max}$ (% FSK) | $24 \pm 2$ | $2 \pm 0$ * | $48 \pm 9$ | $25 \pm 8$ * |
| Log $EC_{50}$ (M) | $-8.7 \pm 0.2$ | $-8.1 \pm 0.3$ | $-8.6 \pm 0.2$ | $-8.5 \pm 0.2$ |
| Hill slope | $0.9 \pm 0.0$ | $0.9 \pm 0.1$ | $0.8 \pm 0.1$ | $0.9 \pm 0.0$ |

**Table 2.** Effect of β-arrestin deletion on agonist-induced cytoplasmic PKA activation in HEK293 cells. Mean parameter estimates ± SEM from responses depicted in Figure 2B, *n*=5. * p<0.05 by paired t-test comparing wild-type *versus* knockout.

|  | **Wild-type** | **β-arrestin-1/2 KO** |
| --- | --- | --- |
| **SNAP-GLP-1R + GLP-1** |  |  |
| $E_{\max}$ (baseline-corrected AUC) | $3.2 \pm 0.4$ | $4.1 \pm 0.5$ |
| Log $EC_{50}$ (M) | $-10.9 \pm 0.1$ | $-11.5 \pm 0.3$ * |
| **SNAP-GIPR + GIP** |  |  |
| $E_{\max}$ (baseline-corrected AUC) | $2.5 \pm 0.5$ | $1.8 \pm 0.2$ |
| Log $EC_{50}$ (M) | $-10.1 \pm 0.0$ | $-10.9 \pm 0.1$ * |
| **SNAP-GCGR + GCG** |  |  |
| $E_{\max}$ (baseline-corrected AUC) | $3.1 \pm 0.5$ | $3.9 \pm 0.2$ |
| Log $EC_{50}$ (M) | $-10.3 \pm 0.1$ | $-11.0 \pm 0.2$ * |

β-arrestins have been observed on many occasions to facilitate signalling by MAPKs such as ERK1/2 (38), and this pathway has been implicated in GLP-1R action in beta cells (30, 39). Interestingly, the peak of ERK1/2 phosphorylation observed after 5 minutes of ligand stimulation was preserved in β-arrestin knockout cells, but duration of the phosphorylation response was reduced for GLP-1R and GCGR, although not for GIPR (Figure 2C). These results are in keeping with newer reports for other GPCRs indicating that the presence of β-arrestins is not essential for ERK1/2 signalling *per se* (36, 40), but may be required for sustained ERK1/2 phosphorylation (39).

To determine the β-arrestin-dependency of incretin receptor endocytosis, we performed further high content microscopy internalisation studies. Surface expression of each receptor was similar in each cell type (Figure 2D). After a 30-min stimulation period, a modest numerical reduction in agonist-induced GLP-1R and GIPR internalisation was noted in β-arrestin knockout cells compared to wild-type, but this was only statistically significant for GIPR. No internalisation of GCGR was detected, as expected. Thus, these experiments corroborate our earlier observations that the absence of β-arrestins has a relatively minor effect on GLP-1R endocytosis (15, 31), with a partial effect seen also with GIPR.

Thus, these results provide initial evidence that β-arrestin recruitment controls both duration and amplitude of cAMP signalling by glucagon family receptors, but is not essential for transient ERK1/2 signalling responses or endocytosis.

### Effects of N-terminal region mutations to GLP-1, GIP and GCG on signalling and trafficking responses

Biased signalling, in which agonists preferentially stabilise certain receptor conformations to engage specific intracellular signalling pathways, may provide a means to selectively increase therapeutic efficacy (41). As the ligand N-terminal region plays a key role in activation of class B GPCRs (42) and is linked to GLP-1R biased signalling (15), we introduced single amino acid substitutions at or close to the N-termini of each endogenous ligand (Table 3 for full amino acid sequences) and tested for intracellular cAMP production (Figure 3A) and β-arrestin-2 recruitment (Figure 3B). For each receptor target, all N-terminally modified analogues retained full efficacy for cAMP, but with reduced potency (full parameter estimates are given in Table 4). Potency for β-arrestin-2 recruitment was also reduced, and in the majority of cases, a reduction in efficacy was also observed. Transduction ratios (43) were calculated to quantify the relative signalling impact of each N-terminal region substitution in each ligand for each pathway (Figure 3C, 3D). Chiral substitution of the first amino acid to dHis1 (GLP-1, GCG) or dTyr1 (GIP), as well as Gly2 and dGln3, were less well tolerated by GIP and GCG than by GLP-1; for example, GIP-dGln3 showed a 100-fold reduction in cAMP potency compared to wild-type GIP, *versus* a 10-fold reduction seen for GLP-1-dGln3 compared to GLP-1. Moreover, comparison of the relative impact of each substitution on cAMP *versus* β-arrestin-2 responses showed that all compounds tested exhibited at least a trend for bias in favour of G protein-dependent cAMP signalling, albeit not statistically significant (as indicated by 95%confidence intervals crossing zero) in some cases (Figure 3E). The large error bars for the bias estimate for GIP-dGln3 reflect the limitations of this method for bias calculation for extremely weak partial agonists (44).

**Table 3.**
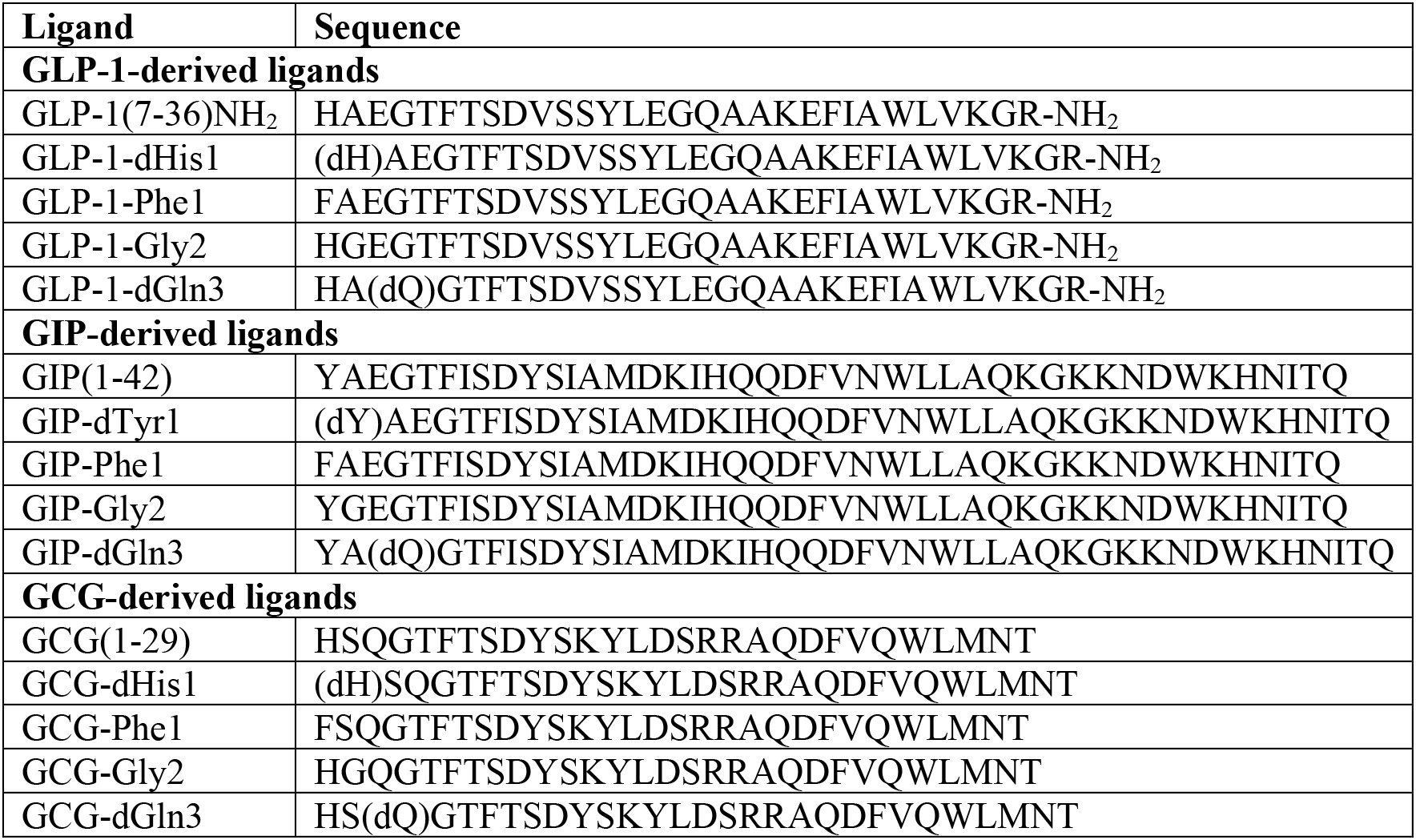
Amino acid sequences of ligands in this study. Sequences are given in standard single letter amino acid code, with D-histidine, D-tyrosine and D-glutamine indicated as “dH”, “dY” and “dQ”, respectively.

**Figure 3.**
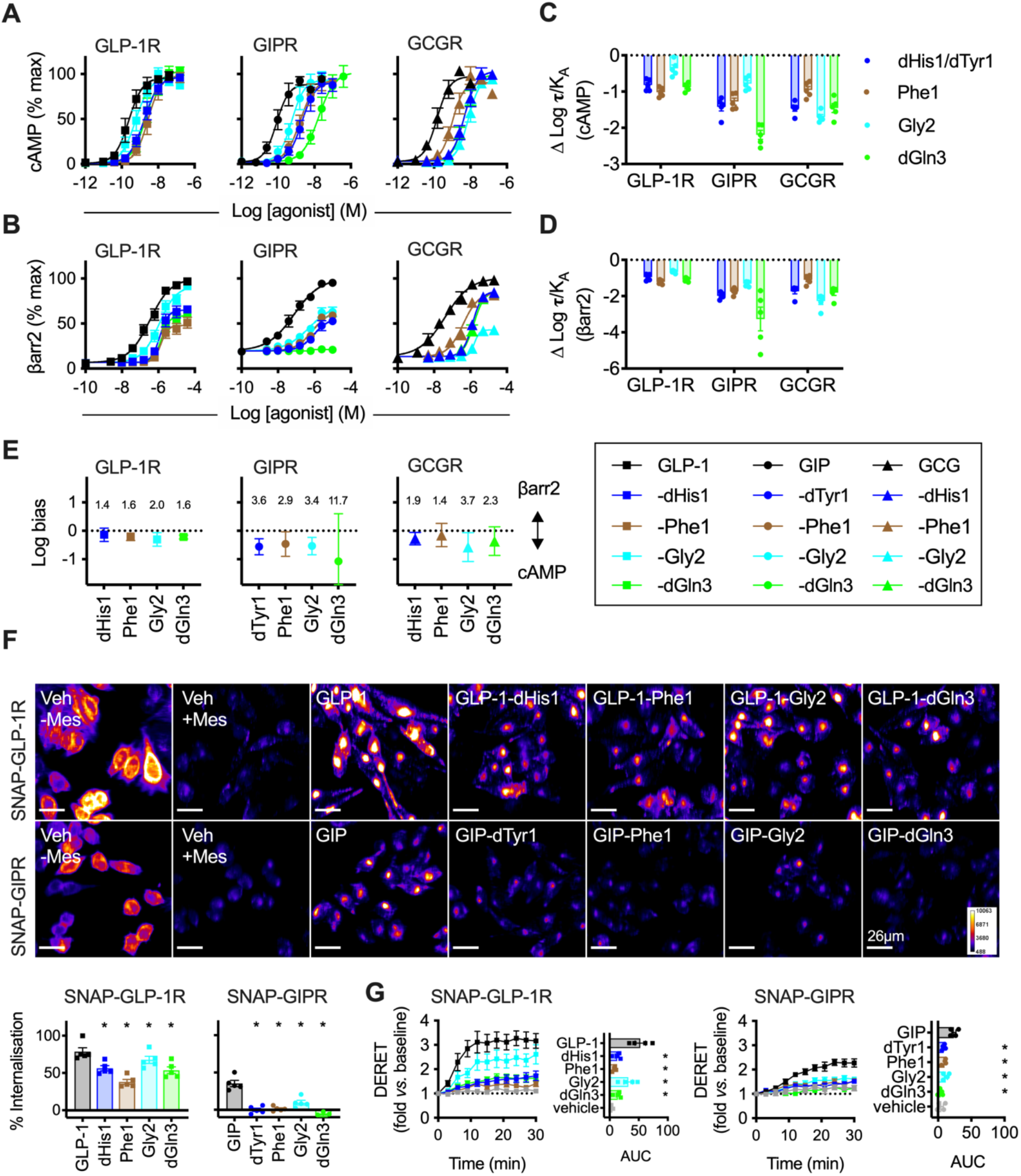
Biased incretin analogues with N-terminal region substitutions. (**A**) cAMP responses in PathHunter CHO-GLP-1R, CHO-GIPR or CHO-GCGR cells (as appropriate) to analogues of GLP-1, GIP and GCG, 30-minute stimulation, *n*=5, 4-parameter fits shown. (**B**) As for (A) but β-arrestin-2 recruitment responses. (**C**) The relative impact of each amino acid substitution on cAMP signalling is shown by subtracting Log **τ**/K_A_ for the reference agonist (GLP-1, GIP or GCG) from that of each analogue on a per-assay basis. (**D**) As for (C) but for β-arrestin-2 recruitment. (**E**) Signal bias for N-terminally modified GLP-1, GIP and GCG analogues at their cognate receptors, calculated as normalised log transduction ratios [ΔΔlog(τ/K_A_)] relative to GLP-1, GIP or GCG, respectively. The numerical degree of bias is indicated for each ligand after anti-log transformation. (**F**) Representative images showing GLP-1R and GIPR internalisation after 30 min stimulation with indicated agonist at 100 nM, with quantification below from *n*=5 experiments and comparison by one-way randomised block ANOVA with Dunnett’s test *versus* GLP-1 or GIP (as appropriate); scale bar = 26 μm. (**G**) SNAP-GLP-1R or SNAP-GIPR internalisation in CHO-K1 cells treated with indicated agonist (100 nM), measured by DERET, *n*=4, AUC *versus* GLP-1 or GIP compared by randomised block one-way ANOVA with Dunnett’s test. * p<0.05 by statistical test indicated in the text. Data are represented as mean ± SEM (with individual replicates in some cases), except for bias plots where error bars indicate 95% confidence intervals.

**Table 4.** Pharmacological characterisation of biased GLP-1R, GIPR and GCGR agonists. Mean parameter estimates ± SEM from responses depicted in Figure 3. * p<0.05, by one-way randomised block ANOVA with Dunnett’s test *vs.* GLP-1, GIP and GCG, as appropriate. Note that for E_max_, statistical comparison was performed prior to normalisation.

| | cAMP | | | | | βarr2 | | | | |
| --- | --- | --- | --- | --- | --- | --- | --- | --- | --- | --- |
| GLP-1R | GLP-1 | dHis1 | Phe1 | Gly2 | dGln3 | GLP-1 | dHis1 | Phe1 | Gly2 | dGln3 |
| $E_{\max}$ (% GLP-1) | 100 | 101 ± 4 | 100 ± 4 | 95 ± 4 | 106 ± 5 | 100 | 66 ± 5 * | 48 ± 6 * | 93 ± 3 | 60 ± 4 * |
| Log $EC_{50}$ (M) | -9.5 ± 0.2 | -8.7 ± 0.2 * | -8.5 ± 0.2 * | -9.2 ± 0.2 * | -8.6 ± 0.2 * | -6.5 ± 0.2 | -5.9 ± 0.1 * | -5.8 ± 0.0 * | -5.9 ± 0.1 * | -5.8 ± 0.0 * |
| Hill slope | 1.7 ± 0.4 | 1.4 ± 0.3 | 1.4 ± 0.1 | 1.4 ± 0.1 | 1.7 ± 0.5 | 0.8 ± 0.1 | 2.0 ± 0.2 * | 2.1 ± 0.2 * | 1.0 ± 0.1 | 2.0 ± 0.3 * |
| Log ($\tau/K_A$) | 9.5 ± 0.2 | 8.7 ± 0.1 * | 8.5 ± 0.2 * | 9.2 ± 0.2 * | 8.6 ± 0.2 * | 6.5 ± 0.1 * | 5.6 ± 0.1 * | 5.3 ± 0.1 * | 5.9 ± 0.1 * | 5.4 ± 0.1 * |
| GIPR | GIP | dTyr1 | Phe1 | Gly2 | dGln3 | GIP | dTyr1 | Phe1 | Gly2 | dGln3 |
| $E_{\max}$ (% GIP) | 100 | 96 ± 6 | 99 ± 3 | 100 ± 4 | 110 ± 4 | 100 | 52 ± 6 * | 65 ± 3 * | 66 ± 7 * | 23 ± 1 * |
| Log $EC_{50}$ (M) | -10.0 ± 0.2 | -8.5 ± 0.2 * | -8.7 ± 0.2 * | -9.1 ± 0.2 * | -7.6 ± 0.3 * | -7.1 ± 0.1 | -6.3 ± 0.2 * | -6.0 ± 0.1 * | -6.5 ± 0.3 | -6.5 ± 0.3 |
| Hill slope | 1.4 ± 0.1 | 1.3 ± 0.2 | 1.2 ± 0.1 | 1.6 ± 0.4 | 1.0 ± 0.1 | 0.6 ± 0.1 | 1.1 ± 0.1 | 1.2 ± 0.2 | 1.0 ± 0.2 | 85.7 ± 53.1 |
| Log ($\tau/K_A$) | 9.9 ± 0.2 | 8.5 ± 0.3 * | 8.6 ± 0.2 * | 9.1 ± 0.2 * | 7.7 ± 0.3 * | 7.2 ± 0.1 | 5.2 ± 0.2 * | 5.5 ± 0.1 * | 5.9 ± 0.1 * | 4.0 ± 0.6 * |
| GCGR | GCG | dHis1 | Phe1 | Gly2 | dGln3 | GCG | dHis1 | Phe1 | Gly2 | dGln3 |
| $E_{\max}$ (% GCG) | 100 | 108 ± 5 | 93 ± 2 | 99 ± 5 | 101 ± 5 | 100 | 85 ± 4 * | 84 ± 4 * | 43 ± 4 * | 85 ± 4 * |
| Log $EC_{50}$ (M) | -9.8 ± 0.2 | -8.2 ± 0.2 * | 8.9 ± 0.2 * | -8.1 ± 0.1 * | -8.3 ± 0.1 * | -7.4 ± 0.3 | -5.9 ± 0.1 * | -6.4 ± 0.1 * | -5.8 ± 0.1 * | -5.8 ± 0.1 * |
| Hill slope | 1.4 ± 0.1 | 1.4 ± 0.2 | 1.6 ± 0.1 | 1.7 ± 0.2 | 1.5 ± 0.1 | 0.6 ± 0.1 | 1.8 ± 0.2 * | 1.7 ± 0.5 * | 2.9 ± 0.4 * | 1.8 ± 0.2 * |
| Log ($\tau/K_A$) | 9.8 ± 0.2 | 8.3 ± 0.1 * | 8.8 ± 0.2 * | 8.1 ± 0.1 * | 8.3 ± 0.1 * | 7.6 ± 0.2 | 5.8 ± 0.1 * | 6.5 ± 0.1 * | 5.3 ± 0.1 * | 5.8 ± 0.1 * |

We also compared GLP-1R and GIPR internalisation induced by these N-terminally substituted agonists, finding that in all cases receptor internalisation was reduced when measured by high content microscopy (Figure 3F) or DERET (Figure 3G) in CHO-K1 cells stably expressing SNAP-GLP-1R or SNAP-GIPR. The GCG analogues were not studied with this assay as GCG itself was without effect.

Overall, these results highlight how the N-termini of each ligand play important roles in receptor activation and initiation of endocytosis. It should be noted that the magnitude of signal bias with the GLP-1 analogues tested here is smaller than for exendin-4-derived biased GLP-1R agonists, a finding that is consistent with our recent exploration of GLP-1/ exendin-4 chimeric peptides (16).

### Biased agonist responses in a beta cell and hepatocyte context

As pancreatic beta cells are a target for GLP-1, GIP and GCG (45), we investigated whether the signalling and trafficking characteristics of N-terminally modified agonists described above could enhance insulin secretion, as previously demonstrated for biased exendin-4-derived GLP-1RAs (15). We used incretin-responsive rat insulinoma-derived INS-1 832/3 cells (46), in which we first confirmed expression of GLP-1R, GIPR and GCGR by qPCR (Figure 4A). Using N-terminally substituted GLP-1 and GIP analogues, we found subtly increased maximal sustained insulin secretion with a number of analogues compared to their respective parent ligand, but this was only apparent at concentrations above 1 μM (Figure 4B, 4C, Table 5). The prolonged incubation in these experiments was specifically selected with the aim of better replicating the *in vivo* situation, where therapeutic ligands with extended pharmacokinetics lead to a state of continuous receptor activation. The majority of analogues displayed reduced potency for insulin secretion, as they had for acute cAMP production in the signalling assays presented in Figure 3. For GIP analogues, maximal insulin secretion was inversely correlated with maximal β-arrestin-2 recruitment, whereas for GLP-1 analogues the relationship was less clear, and in both cases the regression line was shallow (Figure 4D). As GCG can cross-react with GLP-1R in beta cells (47), we tested each N-terminally modified GCG analogue in both wild-type INS-1 832/3 cells and a sub-clone in which GLP-1R had been knocked out by CRISPR/Cas9 (48). This showed that the overall response was dominated by GLP-1R-dependent high dose effects absent in GLP-1R knockout cells, with no clear GCGR-dependent advantageous effect for any analogue (Figure 4E, Table 5).

**Figure 4.**
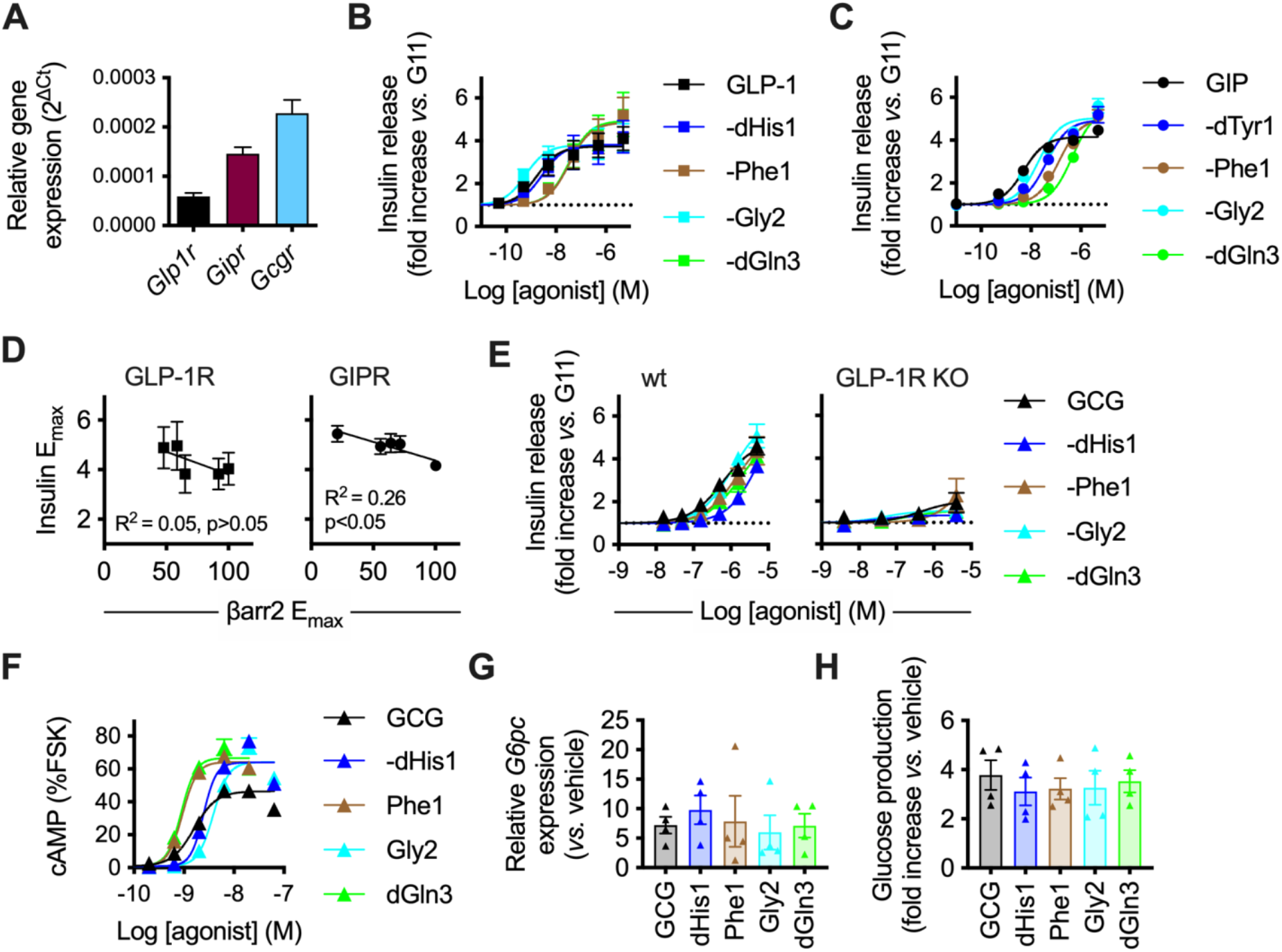
Responses in INS-1 832/3 cells, Huh7 cells and primary hepatocytes. (**A**) Expression of *Glp1r*, *Gipr* and *Gcgr* in INS-1 832/3 cells, determined by qPCR, normalised to expression of endogenous control gene 18S by 2^−ΔCt^, *n*=2. (**B**) Insulin secretory responses in INS-1 832/3 cells treated for 16 hours with GLP-1 analogues at 11 mM glucose (“G11”), *n*=5, 3-parameter fits shown. (**C**) As for (B) but with GIP analogues. (**D**) Correlation of GLP-1 and GIP analogue maximum insulin secretion and maximum β-arrestin-2 recruitment (Figure 3) by linear regression. (**E**) Insulin secretory responses in wild-type and GLP-1R KO INS-1 832/3 cells treated for 16 hours with GCG analogues at 11 mM glucose, *n*=5, 3-parameter fits shown. (**F**) cAMP responses to GCG analogues in Huh7-GCGR cells treated for 16 hours, relative to response to forskolin (10 min, 10 μM), *n*=4, 4-parameter fits shown. (**G**) Effect of prolonged (16-hour) exposure to indicated agonist (10 nM) on upregulation of *G6pc* in Huh7-GCGR cells, *n*=4. (**H**) Effect of prolonged (16-hour) exposure to indicated agonist (100 nM) on glucose production by primary mouse hepatocytes, *n*=4, expressed as fold change to vehicle stimulation. For (G) and (H), no treatment response was significantly different to that of GCG, by one-way randomised block ANOVA with Dunnett’s test. Data are represented as mean ± SEM and individual replicates in some cases.

**Table 5.** Responses to N-terminally substituted ligands in INS-1 832/3 and Huh7-GCGR cells. Mean parameter estimates ± SEM from insulin secretory responses depicted in Figure 4B, 4C and 4E, and 16-hour cAMP accumulation in Figure 4F. Note that the Hill slopes for GCG analogues in Huh7-GCGR cells are derived from the curves plotted from the pooled data, so no statistical comparisons are shown. * p<0.05, by one-way randomised block ANOVA with Dunnett’s test *vs.* GLP-1, GIP and GCG, as appropriate. “n.c.” indicates not calculated.

|  | **GLP-1 analogues - wild-type INS-1 832/3 cells** |  |  |  |  |
| --- | --- | --- | --- | --- | --- |
|  | **GLP-1** | **dHis1** | **Phe1** | **Gly2** | **dGln3** |
| $E_{\max}$ | $4.0 \pm 0.7$ | $3.8 \pm 0.8$ | $4.9 \pm 0.8^*$ | $3.8 \pm 0.6$ | $5.0 \pm 1.0^*$ |
| Log $EC_{50}$ (M) | $-8.5 \pm 0.1$ | $-8.6 \pm 0.0$ | $-7.4 \pm 0.1^*$ | $-9.2 \pm 0.1^*$ | $-7.5 \pm 0.1^*$ |
|  | **GIP analogues - wild-type INS-1 832/3 cells** |  |  |  |  |
|  | **GIP** | **dTyr1** | **Phe1** | **Gly2** | **dGln3** |
| $E_{\max}$ | $4.2 \pm 0.2$ | $4.9 \pm 0.3^*$ | $5.1 \pm 0.4^*$ | $5.0 \pm 0.3^*$ | $5.4 \pm 0.3^*$ |
| Log $EC_{50}$ (M) | $-8.3 \pm 0.1$ | $-7.4 \pm 0.1^*$ | $-6.9 \pm 0.2^*$ | $-7.8 \pm 0.1^*$ | $-6.4 \pm 0.1^*$ |
|  | **GCG analogues – wild-type INS-1 832/3 cells** |  |  |  |  |
|  | **GCG** | **dHis1** | **Phe1** | **Gly2** | **dGln3** |
| $E_{\max}$ | n.c | n.c | n.c | n.c | n.c |
| Log $EC_{50}$ (M) | n.c | n.c | n.c | n.c | n.c |
|  | **GCG analogues – GLP-1R KO INS-1 832/3 cells** |  |  |  |  |
|  | **GCG** | **dHis1** | **Phe1** | **Gly2** | **dGln3** |
| $E_{\max}$ | $1.3 \pm 0.1$ | $1.4 \pm 0.1$ | $1.5 \pm 0.2$ | $1.6 \pm 0.2$ | $1.5 \pm 0.2$ |
| Log $EC_{50}$ (M) | $-7.2 \pm 0.5$ | $-7.3 \pm 0.3$ | $-7.0 \pm 0.8$ | $-6.9 \pm 0.4$ | $-6.5 \pm 0.5$ |
|  | **GCG analogues – Huh7-GCGR cells** |  |  |  |  |
|  | **GCG** | **dHis1** | **Phe1** | **Gly2** | **dGln3** |
| $E_{\max}$ | $46 \pm 1$ | $64 \pm 2^*$ | $64 \pm 4^*$ | $64 \pm 4^*$ | $67 \pm 4^*$ |
| Log $EC_{50}$ (M) | $-8.8 \pm 0.0$ | $-8.6 \pm 0.0^*$ | $-9.0 \pm 0.1^*$ | $-8.4 \pm 0.0^*$ | $-9.1 \pm 0.0^*$ |
| Hill slope | 1.9 | 3.4 | 2.8 | 3.0 | 3.0 |

GCG stimulates glycogenolysis and gluconeogenesis in hepatocytes. Recently, ablation of β-arrestin-2 in hepatocytes was found to increase hepatic glucose output in response to GCG (32). We used Huh7 cells stably expressing GCGR (“Huh7-GCGR”) (49) to assess responses to prolonged stimulation with biased GCG analogues, to determine if differences in β-arrestin recruitment could affect sustained GCGR signalling in a hepatocyte context. Comparisons of maximum cAMP accumulation after 16-hour stimulation with each ligand revealed that greater efficacy for cAMP production was achieved by ligands with reduced β-arrestin-2 recruitment (Figure 4F, Table 5); and in the case of the -Phe1 and -dGln3 compounds, cAMP potency was also greater than for GCG, despite being at least 10-fold less potent acutely (see Figure 3). However, this did not translate to differential changes in GCGR-induced upregulation of the gluconeogenic enzyme glucose-6-phosphatase (G6P; Figure 4G); neither were any significant differences seen for production of glucose in primary mouse hepatocytes (Figure 4H).

Overall, these results indicate that analogues of GLP-1 and GIP with reduced β-arrestin-2 recruitment can augment glucose-stimulated insulin secretion, but with the peptides evaluated in this study, this effect was only apparent at high agonist concentrations. For GCG analogues, prolonged cAMP signalling was seen with agonists displaying reduced β-arrestin-2 recruitment, but this did not translate to increases in downstream responses linked to hyperglycaemia in the models tested.

## Discussion

This study builds on our earlier work using biased GLP-1R agonists derived from exendin-4 and GLP-1 bearing amino acid substitutions close to the N-terminus, in which we also demonstrated prolonged GLP-1R signalling in the absence of β-arrestins but a minor effect of these on endocytosis (15, 16, 31). In the present work we extend these observations to GIPR and GCGR, members of the same class B GPCR family and major investigational targets for metabolic disease. The key findings of this study are: 1) the absence of β-arrestins appears to facilitate prolonged cAMP/PKA signalling with each receptor, with either non-significant or partial effects on endocytosis, and 2) amino acid substitutions at or close to the N-termini of GLP-1, GIP and GCG can diminish β-arrestin-2 recruitment efficacy, with a somewhat lesser effect on cAMP signalling, but the degree of pathway selectivity is reduced compared to what we have previously observed with exendin-4 analogues at the GLP-1R (15), and the impact on prolonged insulin secretion in pancreatic beta cells is more limited.

As expected, GLP-1R, GIPR and GCGR were able to recruit mini-G_s_ and β-arrestin-2 when stimulated with a high concentration of their cognate agonist. However, GIPR responses were in general of reduced amplitude, matching previous observations (24). Interestingly, β-arrestin-2 activation measured by a conformational BRET-based biosensor (27) appeared similar for each receptor, in spite of the difference in recruitment. As proximity-based techniques such as nanoBiT complementation are susceptible to distance constraints imposed by the conformation of the target protein, it is plausible that GIPR-induced β-arrestin-2 recruitment is actually higher than suggested by our results. This might be resolved using a target-agnostic technique such as bystander BRET (50) to monitor β-arrestin recruitment to the plasma membrane, although apparent differences in the segregation of GIPR and GLP-1R into liquid ordered *versus* liquid disordered nanodomains (31) may lead to further confounding depending on acceptor localisation. Further possibilities that could complicate comparisons between the activation and recruitment assays are differences in receptor-effector stoichiometry related to promoter activity, and assay dynamic range.

β-arrestin recruitment to GPCRs sterically blocks G_s_ signalling. Indeed, our data indicate how the absence of β-arrestins prevents a decline in cAMP production during continual agonist stimulation after an initial peak, similar to observations with the β2-adrenergic receptor (36) and adding to the evidence that β-arrestins restrain cAMP signalling at GLP-1R (15) and GCGR (32). Importantly, assuming prolonged activation of each receptor is considered therapeutically desirable, this provides a strong rationale for developing of G protein-biased agonists capable of generating longer lasting signalling responses. We note also that the relative augmentation of cAMP production at the later time-point in β-arrestin knockout cells was similar for each receptor, in keeping with their apparently similar ability to induce β-arrestin-2 conformational change (as in Figure 1). Contrasting with the situation for cAMP/PKA, agonist-induced ERK1/2 phosphorylation tended to be preserved at early time-points in β-arrestin knockout cells but reduced at later time-points. Multiple factors are implicated in incretin receptor-mediated ERK1/2 phosphorylation (42, 51, 52), and for GPCRs more widely there is ongoing controversy concerning the relative contribution of G proteins *versus* β-arrestins to MAPK activation (40, 53). Nevertheless, our data suggests that β-arrestins may indeed play a role specifically in sustained (rather than acute) ERK1/2 signalling, which has been implicated in GLP-1R-induced protection against apoptosis in beta cells (39). Via a different mechanism, ERK1/2 is also implicated in GIPR-mediated beta cell survival (54), although we did not observe reduced GIP-induced ERK1/2 phosphorylation in β-arrestin knockout cells in our experiments. It is unclear whether potential reductions in signalling pathways engaged by ERK1/2 and other putative β-arrestin-scaffolded MAPKs are relevant to the therapeutic action of incretin receptor-targeting biased ligands. Reassuringly, there was no evidence of reduced beta cell survival in mice chronically treated with a biased GLP-1R agonist with undetectable β-arrestin recruitment (55).

Increasing emphasis is now placed on understanding ligand-specific effects on receptor trafficking due to its potential importance in spatiotemporal control of intracellular signalling (9). In our hands, the absence of both β-arrestin isoforms had a surprisingly small effect on GLP-1R, which is in keeping with our earlier data (15, 31). For GIPR, a partial reduction in internalisation was observed, consistent with the report of Gabe *et al*., who showed partially reduced GIPR internalisation in the same cell models (56) using DERET. Overall, both GLP-1R and GIPR can continue to undergo ligand-induced endocytosis in the absence of β-arrestins, suggesting the existence of β-arrestin-independent endocytic mechanism(s). We cannot, however, exclude the possibility of compensatory upregulation or rewiring of secondary endocytic pathways in β-arrestin knockout cells, which could disguise a more significant role for β-arrestins in the endocytosis of these receptors.

GLP-1 is more dependent on its N-terminus for binding to the GLP-1R than is exendin-4 (57). Sequential truncation of the first nine amino acids of exendin-4 results in only a modest reduction in binding affinity, but virtually abolishes binding of GLP-1 (22, 58). Thus, the reduction in signalling potency resulting from an N-terminal amino acid substitution within the GLP-1 backbone may be secondary to reduced affinity, whereas the same change in exendin-4 might have little impact on occupancy, thereby allowing the modified ligand to achieve biased signalling at higher potency. In agreement with this concept, acute cAMP signalling potencies for exendin-dHis1 and exendin-Phe1 in our earlier study were, respectively, no different to and 2.5-fold lower than for exendin-4 (15), while contrastingly in the present work, the same substitutions to the GLP-1 N-terminus reduced cAMP potency by, respectively, a factor of 6 and 10. This might limit the potential for these modified GLP-1 analogues to improve downstream signalling outputs during prolonged stimulation, except at maximal doses when receptor occupancy is high, and even then, reduced mini-G_s_ recruitment is clearly demonstrated with other GLP-1R agonists with N-terminal modifications at supramaximal stimulatory concentrations (16). As for GLP-1, the N-termini of both GCG and GIP are also known to play a major role in affinity for their cognate receptors, with truncation of the terminal amino acid residue resulting in a >10-fold loss of affinity in both cases (59, 60). The resulting reductions in agonism have the potential to partly counterbalance benefits from reduced β-arrestin-mediated desensitisation, partly reconciling the discrepancy between the modest increases in insulinotropic efficacy over 16 hours with N-terminally modified ligands *versus* the striking differences in duration of cAMP signalling in β-arrestin knockout cells over 60 minutes with non-modified GLP-1, GCG and GIP.

A number of analogues tested in this report have previously been described, due in part to the interest in reducing ligand sensitivity to degradation by the N-terminal targeting exopeptidase DPP-4 (61). Published potency or affinity measures for GLP-1-dHis1 (62) and - Gly2 (63) were broadly in agreement with our results, although GLP-1-Phe1 was found to be well tolerated for cAMP signalling potency in RIN-T3 cells (64), contrasting with the deleterious effect we observed. Differences in cell type, receptor species, incubation times and other factors may influence responses to agonists, complicating direct comparisons with reported values in the literature (65). The affinity of GIP-dTyr1 was reduced 10-fold compared to unmodified GIP (66), similarly to our results, whereas the -Gly2 substitution was well tolerated (67). GIP-Phe1 has been used as a GIPR I^125^-radioligand (68). These datasets are complemented here by our measures of bias between cAMP and β-arrestin recruitment, and endocytosis, which have not previously been reported for these ligands, or indeed for any putative biased GCGR or GIPR agonist to our knowledge.

A further factor which might contribute to the relative lack of effect on downstream responses to biased GLP-1, GIP and GCG analogues during prolonged incubations is enzymatic peptide degradation, for example by neutral endopeptidase 24.11 (NEP 24.11), found on pancreatic beta cell membranes and capable of hydrolysing GLP-1, GCG, and to a lesser extent, GIP (69), or endothelin converting enzyme-1 (70) situated predominantly in endosomal compartments. DPP-4, also expressed by beta cells (71), is also likely to contribute, although the modified N-termini of the ligands tested may confer some resistance to its action. Sequence optimisation to increase proteolytic stability during our extended *in vitro* studies may be required to maintain adequate ligand concentration to fully manifest consequences of signal bias. In the *in vivo* setting, fatty acid conjugation such as in liraglutide (72) protects against NEP 24.11 and DPP-4 degradation, presumably as the resultant albumin-bound form of the ligand is inaccessible to the enzymes. One possible future approach would be to test acylated forms of the ligands described herein to determine if sustained exposure to the N-terminally substituted forms led to enhanced metabolic effects. Additionally, as our beta cell studies were performed with INS-1 832/3 clonal beta cells, it would be important to validate key findings in primary islets, ideally from humans, to ensure observations are not an artefact of the model used.

Whilst GLP-1R agonists developed specifically to G protein-directed signalling are yet to be tested in humans, the potential utility of this approach is supported by the recent observation that Tirzepatide, a dual GLP-1R/GIPR agonist peptide currently in late stage clinical trials (17), and its non-acylated precursor show a significant degree of signal bias for the GLP-1R in favour of cAMP over β-arrestin recruitment (18, 73). Conflicting reports exist for signal bias between cAMP and β-arrestin recruitment to the GIPR for Tirzepatide, with one study showing bias in favour of cAMP and another showing no difference (18, 74). Signal bias at the GCGR is relatively unexplored, except for a recent study of a series of dual GLP-1R/GCGR agonists in which small response amplitude for β-arrestin-2 recruitment to GCGR hampered bias assessments (75), but should be further explored in the future.

In summary, we demonstrate in this study that GLP-1, GIP and GCG analogues with a variety of N-terminal substitutions typically show reduced β-arrestin-2 recruitment. In the case of GLP-1 and GIP, this is associated with reduced receptor endocytosis, and this effect can be exploited to increase maximal insulin release *in vitro*. Generation of long-lasting biased incretin mimetics will be required to determine whether this applies *in vivo*.

## Experimental procedures

### Peptides

All peptides were obtained from Insight Biotechnology and were certified by HPLC to be at least 90% pure.

### Cell lines

HEK293T cells were maintained in DMEM, 10% FBS and 1% penicillin/streptomycin. Wild-type and dual β-arrestin knockout (36) HEK293 cells were maintained in DMEM, 10% FBS and 1% penicillin/streptomycin. Monoclonal CHO-K1 cells stably expressing SNAP-GLP-1R or SNAP-GIPR (31) were maintained in Ham’s F12 medium, 10% FBS and 1% penicillin/streptomycin. PathHunter β-arrestin-2 CHO-K1 cells (DiscoverX) stably expressing human GLP-1R, GIPR, or GCGR were maintained in the manufacturer’s proprietary culture medium. INS-1 832/3 cells (46), a gift from Professor Christopher Newgard, were maintained in RPMI supplemented with 11 mM glucose, 10% FBS, 10 mM HEPES, 2 mM L-glutamine, 1 mM pyruvate, 50 μM β-mercaptoethanol and 1% penicillin/streptomycin. INS-1 832/3 cells lacking endogenous GLP-1R after deletion by CRISPR/Cas9 (48), a gift from Dr Jacqueline Naylor, AstraZeneca, were maintained similarly. A stable clone of Huh7 hepatoma cells expressing human GCGR was generated from a previously described multi-clonal cell population (49) by flow cytometric sorting of cells labelled with FITC-conjugated glucagon, and subsequently maintained in DMEM, 10% FBS, 1% penicillin/streptomycin and 1 mg/ml G418.

### Isolation of primary hepatocytes

Hepatocytes from adult male C57Bl/6J mice were isolated using collagenase perfusion (76). After filtering and washing, cells were plated in 12-well collagen-coated plates at 3× 10^5^ cells/ml, 1 ml per well in attachment medium (M199 with 1% penicillin/streptomycin, 1% BSA, 10% FBS, 100 nM triiodothyronine, 100 nM dexamethasone and 100 nM insulin). After 5 hours, attachment medium was replaced with serum starvation medium (M199 with 1% penicillin/streptomycin, 100 nM dexamethasone and 10 nM insulin).

### Measurement of mini-G and β-arrestin-2 recruitment by nanoBiT complementation

The plasmids for mini-G_s_, -G_i_ and -G_q_, each tagged at the N-terminus with the LgBiT tag (20), were a kind gift from Prof Nevin Lambert, Medical College of Georgia. The plasmid for β-arrestin-2 fused at the N-terminus to LgBiT was obtained from Promega (plasmid no. CS1603B118). Construction of the GLP-1R-SmBiT plasmid was described previously (55) and the same strategy was used to develop GIPR-SmBiT and GCGR-SmBiT, with cloning in frame at the C-terminus of the receptor by substitution of the Tango sequence on a FLAG-tagged GPCR-Tango expression vector (77), a gift from Dr Bryan Roth, University of North Carolina (Addgene # 66295). HEK293T cells in 12-well plates were co-transfected using Lipofectamine 2000 with the following amounts of plasmid DNA: 0.5 μg of GPCR-SmBiT plus 0.5 μg LgBiT-mini-G_s_, -G_i_ or -G_q_; or 0.05 μg each of GPCR-SmBit and LgBit-β-arrestin-2 plus 0.9 μg empty vector DNA (pcDNA3.1). After 24 hours, cells were resuspended in Nano-Glo dilution buffer + furimazine (Promega) diluted 1:50 and seeded in 96-well half area white plates. Baseline luminescence was measured over 5 minutes using a Flexstation 3 plate reader at 37°C before addition of agonist or vehicle. After agonist addition, luminescent signal was serially recorded over 30 min, and normalised to well baseline and then to average vehicle-induced signal to establish the agonist effect.

### Measurement of β-arrestin-2 activation by intramolecular BRET

HEK293T cells in 6-well plates were co-transfected using Lipofectamine 2000 with the following amounts of plasmid DNA: 0.5 μg SNAP-GLP-1R, SNAP-GIPR or SNAP-GCGR (all from Cisbio), 0.5 μg Nluc-4myc-β-arrestin-2-CyOFP1 (27), and 1 μg pcDNA3.1. After 24 hours, cells were resuspended in Nano-Glo dilution buffer + furimazine (1:50) and seeded in 96-well half area white plates. Baseline luminescence was measured at 460 nm and 575 nm over 5 minutes using a Flexstation 3 plate reader at 37°C before addition of agonist or vehicle. After agonist addition, luminescent signals at the same wavelengths were serially recorded over 15 min. The BRET ratio (575/460) was calculated at each timepoint, normalised to well baseline and then to average vehicle-induced signal to establish the agonist-induced BRET effect.

### Measurement of receptor internalisation by DERET

Diffusion-enhanced resonance energy transfer (DERET) (7) was used to monitor agonist-induced receptor internalisation in HEK293T cells transiently transfected for 24 hours with SNAP-tagged receptors (2 μg plasmid DNA per well of 6-well plate), or in monoclonal CHO-K1 cells stably expressing SNAP-GLP-1R or SNAP-GIPR. Labelling was performed using the time-resolved Förster resonance energy transfer (TR-FRET) SNAP-probe Lumi4-Tb (Cisbio) at 40 nM for 60 minutes at room temperature, either in suspension (for HEK293T) or with adherent cells (for CHO-K1). After washing three times, fluorescein (24 μM in HBSS) was added to cells in opaque bottom white plates, and baseline signal was read for 10 min using a Flexstation 3 plate reader (λ_ex_ 340 nm, λ_em_ 520 and 620 nm, delay 400 μs, integration 1500 μs) at 37°C. Agonists, prepared in 24 μM fluorescein, were added, and signal was sequentially monitored. Receptor endocytosis leads to reduced contact of Lumi4-Tb with extracellular fluorescein, and a reduction in signal at 520 nm with an increase at 620 nm. After first subtracting values from wells containing fluorescein only, internalisation was expressed ratiometrically as signal obtained at 620 nm divided by that obtained at 520 nm.

### Measurement of receptor internalisation using a cleavable SNAP-labelling probe

The assay was adapted from a previous description (16). HEK293T or wild-type / dual β-arrestin knockout HEK293 cells were seeded in black, clear bottom, plates coated with 0.1% poly-D-lysine, and assayed 24 hours after transfection with SNAP-tagged GLP-1R, GIPR or GCGR plasmid DNA (0.1 μg per well). Cells were labelled with the cleavable SNAP-tag probe BG-S-S-649 (featuring the DY-649 fluorophore, a gift from New England Biolabs) in complete medium for 30 minutes at room temperature. After washing, fresh medium ± agonist was added, with agonists applied in reverse time order in the case of time-course experiments. At the end of the incubation, medium was removed and wells were treated with for 10 min at 4°C with Mesna (100 mM, in alkaline TNE buffer, pH 8.6) to remove BG-S-S-649 bound to residual surface receptor without affecting the internalised receptor population, or with alkaline TNE buffer alone. After washing, cells were imaged using an automated Nikon Ti2 widefield microscope with LED light source (CoolLED) and a 0.75 numerical aperture 20X air objective, assisted by custom-written high content analysis software (78) implemented in Micro-Manager (79). A minimum of 4 epifluorescence and matching transmitted phase contrast images per well were acquired. Average internalised receptor across the imaged cell population was quantified using Fiji as follows: 1) phase contrast images were processed using PHANTAST (80) to segment cell-containing regions from background; 2) illumination correction of fluorescence images was performed using BaSiC (81); 3) fluorescence intensity was quantified for cell-containing regions. Agonist-mediated internalisation was determined by comparing the mean signal for each condition normalised to signal from wells not treated with Mesna, after first subtracting non-specific fluorescence determined from wells treated with Mesna but no agonist.

### Visualisation of receptor redistribution

HEK293T cells seeded on 0.1% poly-D-lysine-coated coverslips were transiently transfected for 24 hours with SNAP-GLP-1R, SNAP-GIPR or SNAP-GCGR (0.5 μg per well of 24-well plate). Surface SNAP-tag labelling was performed using SNAP-Surface-549 (1 μM) for 30 min at 37°C. After washing, cells were stimulated ± 100 nM agonist for 30 min at 37°C, followed by fixation with 2% paraformaldehyde. Coverslips were mounted using Diamond Prolong antifade with DAPI and imaged using a 1.45 numerical aperture 100X oil immersion objective, with z-stacks acquired throughout the cell volume with a step size of 0.2 μm. Deconvolution was performed using Deconvolutionlab2 (82), and a maximum intensity projection from 10 consecutive z-planes was constructed to generate the final images.

### Cyclic AMP assays

All experiments were performed at 37°C. <u>Wild-type and dual β-arrestin knockout HEK293 cells:</u> 24 hours after transient transfection with SNAP-GLP-1R, SNAP-GIPR or SNAP-GCGR (1 μg per well of 12-well plate), cells were resuspended in serum-free Ham’s F12 medium and stimulated with the indicated agonist for 10 or 60 min without phosphodiesterase inhibitors. Forskolin (10 μM) was used as a control. cAMP was quantified by HTRF (cAMP Dynamic 2, Cisbio), and responses were normalised to that of forskolin. <u>PathHunter CHO-K1 cells:</u> Cells were resuspended in serum-free Ham’s F12 medium and treated with indicated agonist, without phosphodiesterase inhibitors, for 30 minutes, followed by application of detection reagents for determination of cAMP by HTRF. β-arrestin-2 recruitment responses were measured in parallel. <u>Huh7-GCGR</u> cells were treated with indicated concentration of agonist without phosphodiesterase inhibitors before lysis. 3- or 4-parameter curve fitting was performed using Prism 8.0 (Graphpad Software).

### Measurement of PKA activation

After co-transfection for 36 hours with plasmid DNA encoding the relevant SNAP-tagged receptor (1 μg) and AKAR4-NES (1 μg; a gift from Dr Jin Zhang, Addgene plasmid #647270), wild-type or dual β-arrestin knockout HEK293 cells were suspended in HBSS in black 96 well plates. After a 10-min baseline measurement, compounds were added and fluorescence measured sequentially using a Flexstation 3 plate reader (λ_ex_ = 440 nm, λ_em_ = 485 nm and 535 nm). After blank well subtraction, signals were expressed ratiometrically and agonist-induced changes calculated relative to individual well baseline. Curve fitting was performed to determine EC_50_ from overall signal AUC.

### Measurement of ERK1/2 phosphorylation

24 hours after transient transfection with SNAP-GLP-1R, SNAP-GIPR or SNAP-GCGR (0.1 μg per well of 96-well plate), wild-type or dual β-arrestin knockout HEK293 cells were stimulated in reverse time order with the indicated ligand (100 nM) in serum-free Ham’s F12 medium. ERK1/2 phosphorylation was determined by HTRF (Cisbio Phospho-ERK [Thr202/Tyr204] cellular kit) from cell lysates prepared using the manufacturer’s lysis buffer with phosphatase/protease inhibitors. Ligand-stimulated HTRF ratios were normalised for each experiment as a fold change of the HTRF ratio from unstimulated cells.

### Measurement of β-arrestin recruitment by enzyme fragment complementation

β-arrestin-2 recruitment was determined by enzyme fragment complementation using the PathHunter system (DiscoverX). CHO-K1 cells expressing GLP-1R, GIPR or GCGR were treated with indicated concentrations of agonist for 30 minutes at 37°C before addition of detection reagents and read by luminescence.

### Gene expression analysis

RNA was harvested from INS-1 832/3 and Huh7 cells using the Cells-to-CT kit (Themo Fisher). Taqman probes were used to detect expression of *Glp1r* (Rn00562406_m1), *Gipr* (Rn00562325_m1), *Gcgr* (Rn00597162_g1), *G6pc* (Rn00689876_m1) and endogenous control gene 18S (Hs99999901_s1) by quantitative PCR.

### Insulin secretion

Insulin secretion from INS-1 832/3 cells (46) was assayed after a prior overnight period of exposure to low glucose (3 mM) complete medium. Cells were added in suspension to plates containing indicated agonists, prepared in RPMI containing 2% FBS and 11 mM glucose, for 16 hours. Supernatant insulin concentration was determined by HTRF (High Range Insulin kit, Cisbio). Results were normalised to those obtained with 11 mM glucose but no additional agonist. Three-parameter fitting was performed using Prism 8.0.

### Data analysis and statistics

All analyses were performed using Prism 8.0. For bias calculations, to reduce contribution of inter-assay variability, cAMP and β-arrestin-2 assays were performed concurrently, with the same incubation time of 30 minutes to avoid artefactual bias resulting from different activation kinetics of each pathway (65); bias was determined by calculating transduction coefficients (43, 83); here, due to the matched design of our experiments, we calculated ΔΔlog(τ/K_A_) on a per-assay basis by normalising the log(τ/K_A_) of each ligand to the relevant endogenous ligand (GLP-1, GIP or GCG, to generate a Δlog[τ/K_A_] value) and then to the reference pathway (cAMP). For experiments with a matched design, paired two-tailed t-tests or randomised block ANOVAs were performed. Specific statistical tests are indicated in the figure legends. Statistical significance was inferred when p<0.05. To determine statistical significance for signal bias, 95% confidence intervals were calculated; bias *versus* the reference endogenous ligand was considered statistically significant when this confidence interval did not cross zero, as previously recommended (83).

### Data availability

Any additional data supporting the analyses in the manuscript is available from B.J. on reasonable request.

## Author contributions

B.J., S.R.B. and A.T. conceived and designed the study. B.J., E.R.M., Z.F., M.L. and P.P. performed and analysed experiments. I.R.C., A.O. and R.J. and A.I. provided novel reagents. S.K., F.G., C.D. and P.M.W.F. developed HCA instrumentation and analysis tools. Funding for the study was acquired by B.J., A.T., S.R.B., G.A.R. and T.T. B.J. wrote the draft manuscript. All authors made manuscript revisions and approved the final version.

## Abbreviations

cAMP: Cyclic adenosine monophosphate
DERET: Diffusion-enhanced resonance energy transfer
DPP-4: Dipeptidyl dipeptidase-4
GCG: Glucagon
GCGR: Glucagon receptor
GIP: Glucose-dependent insulinotropic polypeptide
GIPR: Glucose-dependent insulinotropic polypeptide receptor
GLP-1: Glucagon-like peptide-1
GLP-1R: Glucagon-like peptide-1 receptor
NEP24.11: Neutral endopeptidase
T2D: Type 2 diabetes
TR-FRET: Time-resolved Förster resonance energy transfer

