## Supporting Information for "Signal bias at glucagon family receptors: rationale and downstream impacts"

### Impact of N-terminally substituted glucagon family receptor agonists on signal bias, trafficking and downstream responses

Ben Jones[1,#], Emma Rose McGlone[1], Zijian Fang[1,2], Phil Pickford[1], Ivan R Corrêa Jr[3], Atsuro Oishi[4,5,6,7], Ralf Jockers[4,5,6], Sunil Kumar[8], Frederik Görlitz[8], Chris Dunsby[8], Paul MW French[8], Guy A Rutter[9,10], Tricia Tan[1], Alejandra Tomas[9,#], Stephen R Bloom[1].

[1]Section of Endocrinology and Investigative Medicine, Imperial College London, London, United Kingdom.
[2]Current address: Wellcome Trust – Medical Research Council Cambridge Stem Cell Institute and Department of Haematology, University of Cambridge, Cambridge, United Kingdom
[3]New England Biolabs, Ipswich, USA
[4]Inserm U1016, Institut Cochin, Dept Endocrinology, Metabolism and Diabetes, Paris, France.
[5]CNRS UMR 8104, Paris, France.
[6]University Paris Descartes, Sorbonne Paris Cité, Paris, France.
[7]Current address: Department of Anatomy, Kyorin University Faculty of Medicine, Tokyo, Japan.
[8]Department of Physics, Imperial College London, London, United Kingdom
[9]Section of Cell Biology and Functional Genomics, Imperial College London, London, United Kingdom.
[10]Lee Kong Chian School of Medicine, Nanyang Technological University, Singapore.

**Running title:** Signal bias at glucagon family receptors

#### Supplementary Figure 1

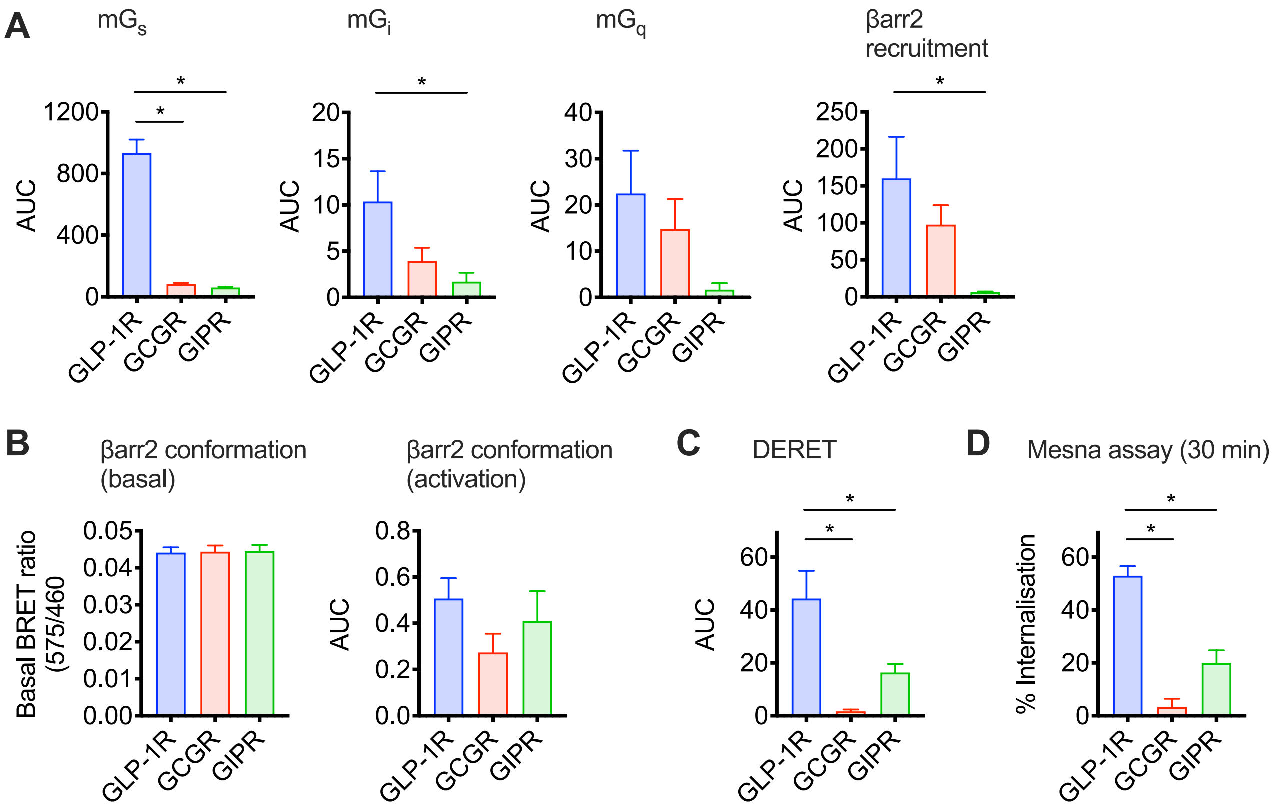


**Supplementary Figure 1. Recruitment and signalling responses.** (**A**) AUC analyses from nanoBiT kinetic responses shown in Figure 1A, with comparisons by one-way ANOVA with Tukey's test. (**B**) Basal (unstimulated) BRET ratio and AUC analysis from β-arrestin-2 activation BRET assay shown in Figure 1B, with comparisons by one-way ANOVA with Tukey's test. (**C**) AUC analysis from DERET assay shown in Figure 1C, with comparisons by randomised block one-way ANOVA with Tukey's test. (**D**) Statistical comparison of internalisation measured by reversible surface SNAP-labelling (see Figure 1D, E) using randomised block one-way ANOVA with Tukey's test. * $p<0.05$ by statistical test indicated in the text. Data are represented as mean ± SEM.

#### Supplementary Figure 2

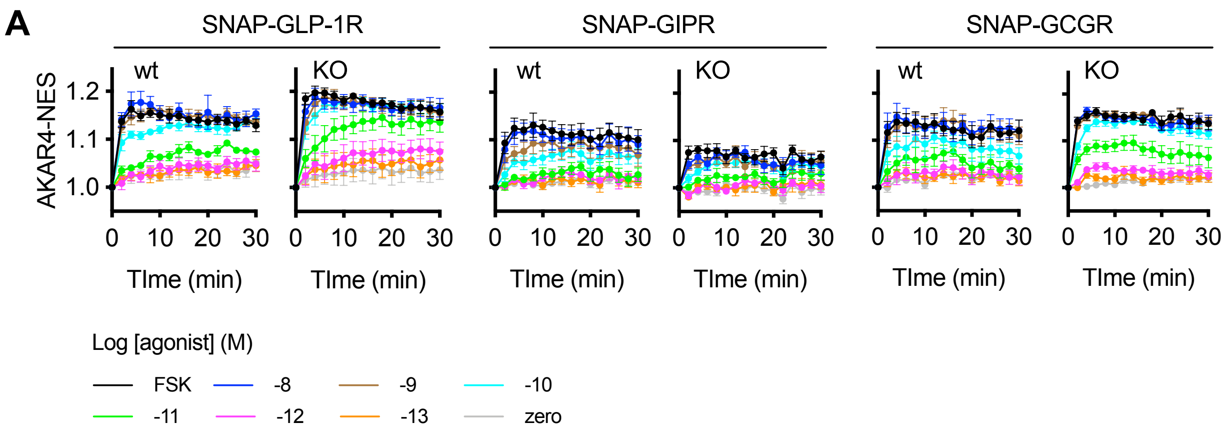


**Supplementary Figure 2. Responses in wild-type *versus* dual β-arrestin knockout cells.** (**A**) Cytoplasmic PKA activation in wild-type or dual β-arrestin knockout HEK293 cells transiently transfected with AKAR4-NES and indicated SNAP-tagged receptor, stimulated with indicated concentration of GLP-1, GIP or GCG, or forskolin (10 µM), with FRET signal indicated ratiometrically after normalisation to individual well baseline, *n*=5. Data are represented as mean ± SEM.
